# Heterobenzamides exhibit bacteriostatic activity against intracellular *Mycobacterium tuberculosis* by targeting aerobic respiration

**DOI:** 10.64898/2026.08.25.747121

**Authors:** Aditi Deshpande, Tanya Parish

**Author notes:** Department of Internal Medicine, UT Southwestern Medical Center, Dallas, TX, United States.

## Abstract

We previously identified a series of heterobenzamides (HBAs) with potent growth inhibitory activity against *Mycobacterium tuberculosis* in axenic culture. We also provided evidence that these target QcrB, a component of the terminal cytochrome oxidase in the electron transport chain. We expanded our studies to look at the full microbiological profile: key molecules from the series were tested for activity under different conditions and against additional strains. HBA analogs were active against intracellular bacteria where they exhibited bacteriostatic activity. A strain of *M. tuberculosis* with a mutation in QcrB (T313I) was resistant to HBAs in both axenic culture and inside macrophages. HBAs retained potency against lineages and mono-resistant strains of *M. tuberculosis*. HBAs had a narrow spectrum of activity, since they were not active against the “ESKAPEE” pathogens. Combination of the key HBA with bedaquiline was synergistic, as expected for a QcrB inhibitor, but there was no strong synergy with other drugs. Exposure of *M. tuberculosis* to the key HBA led to ATP depletion and boosted the oxygen consumption rate. This effect was specific to *M. tuberculosis*, since human THP-1 macrophage-like cells were unaffected by exposure to the HBA. HBA did not induce the production of reactive oxygen species or affect membrane potential but did affect pH homeostasis. Taken together, these data provide further evidence to support the identification of QcrB as the target and indicate that they are suitable for further drug development.

## INTRODUCTION

Tuberculosis (TB), caused by the bacterial pathogen *Mycobacterium tuberculosis* remains one of the oldest global diseases impacting human health. According to the WHO, in 2024, more than 10.5 million people were infected with *M. tuberculosis*, leading to more than 1 million fatalities throughout the world (1). Lengthy anti-TB treatment and the rise in emergence of drug-resistant *M. tuberculosis* strains demands discovery and development of new drugs.

*M. tuberculosis*, an aerobic pathogen, uses oxidative phosphorylation for energy generation (2). Energy production is essential to *M. tuberculosis* under replicating and non-replicating conditions (3). Targeting essential energy-generating pathways has been a productive strategy in the development of new anti-tubercular drugs (4–6).

Aerobic respiration in *M. tuberculosis* utilizes a branched electron transport chain (ETC) terminating in cytochrome oxidases which pump electrons across the cytoplasmic membrane. The proton motive force (PMF) then drives ATP synthesis via the bacterial F0-F1 ATPase. *M. tuberculosis* has two terminal cytochrome oxidases – cytochrome bc and cytochrome bd oxidase (2, 7). Under aerobic conditions, the energetically efficient cytochrome bc is preferred over the less efficient cytochrome bd oxidase. When cytochrome bc is inhibited, electrons are diverted via cytochrome bd oxidase. Since *M. tuberculosis* has an alternative electron acceptor, cytochrome bc inhibitors are generally bacteriostatic extracellularly, in standard medium. However, during infection, cytochrome bc function is essential, since Q203 (Telacebec), an inhibitor of the QcrB subunit. has demonstrated activity in human clinical trials and thereby provides a clinical validation of QcrB as a target (7, 8).

In addition to the Q203 series, numerous other series with different chemical scaffolds and physicochemical properties have been identified which target QcrB and have similar microbiological profiles (7, 9–15). Among these, we previously identified the heterobenzamides series with good activity in aerobic culture and provided confirmation of on-target activity (against QcrB) (9).

Here, we expand our previous work to characterize the activity of HBA analogs under various conditions and provide further evidence that these molecules target energy generation via disruption of the electron transport chain.

## METHODS

### Bacterial and cell culture

*M. tuberculosis* strains were cultured in Middlebrook 7H9 medium supplemented with 10 % v/v Middlebrook OADC supplement and 0.05 % w/v Tween 80 (7H9-OADC-Tw) at 37 °C unless otherwise stated. *M. tuberculosis* strains constitutively expressing luciferase from the pMV306hsp+LuxG13 plasmid were grown in 7H9-OADC-Tw medium supplemented with 20 µg/mL kanamycin.(16) 7H9-OADC-Tw medium supplemented with 40 mM sodium pyruvate was used to grow *M. tuberculosis* strains N1201 (Lineage 5) and N1268 (Lineage 6). *M. tuberculosis* H37Rv-LP (ATCC 25618) expressing a pH-responsive ratiometric GFP (rGFP)(17) was cultured in 7H9-OADC-Tw medium supplemented with 50 µg/mL hygromycin. *Escherichia coli* ATCC BW25113, *Staphylococcus aureus* ATCC 12600, *Klebsiella pneumoniae* ATCC 13883, *Acinetobacter baumannii* ATCC 19606, *Pseudomonas aeruginosa* ATCC 10145, *Enterobacter cloacae* ATCC 13047 were cultured in Cation-adjusted Mueller Hinton Broth (CAMHB). *Enterococcus faecium* ATCC 19434 was cultured in Brain heart infusion (BHI) broth. THP-1 cells obtained from ATCC (TIB-202) were cultured in cRPMI 1640 or complete RPMI 1640 (RPMI 1640 supplemented with 10% v/v fetal bovine serum and 10% v/v Glutagro) and incubated at 37°C with 5% CO_2_.

### Determination of minimum inhibitory concentration (MIC)

*M. tuberculosis* strains were cultured to mid-logarithmic phase (OD∼0.4-0.6) and inoculated to a final OD of 0.02 in 96-well plates containing 10-point, 2-fold dilutions of the test compounds. Growth was measured by OD_590_ using a Synergy H4 plate reader after 5 days for *M. tuberculosis* H37Rv-LP and QcrB_T313I_ strains and 7 days for all other strains. Viability was measured after the addition of Alamar blue for 24 h for strains N1201 and N1268 and measured by fluorescence at Ex560/Em590 nm. *Escherichia coli* , *Enterobacter cloacae, Staphylococcus aureus, Klebsiella pneumoniae, Acinetobacter baumannii* , *Pseudomonas aeruginosa*, and *Enterococcus faecium* were cultured for 18 hours, diluted, and inoculated into 96-well plates at a final OD of 0.04 for *E. coli*, 0.02 for *P. aeruginosa* and *E. cloacae*, 0.01 for *K. pneumoniae* and *S. aureus*, 0.001 for *E. faecium*. Assay plates were incubated for 18 h at 37°C and OD_590_ was read using a Synergy H1 plate reader. IC_90_ (the concentrations at which 90% growth was inhibited) values were calculated by fitting dose-response curves using the variable Hill slope model.

### Determination of intracellular and bacteriostatic activity

THP-1 cells were differentiated with 80 nM phorbol 12-myristate-13 acetate (PMA) overnight and infected with *M. tuberculosis* at a multiplicity of infection (MOI) of 1:1. THP-1 cells were recovered using Accumax™, incubated at RT for 20 min, harvested and resuspended in fresh complete RPMI (cRPMI) medium. Cells were inoculated into 96-well plates at a final density of 4×10^5^ cells/mL and incubated at 37°C and 5% CO_2._ At time 0, one set of plates was treated with 25 µL Accumax™ at RT for 10 min; samples were diluted 1:10 in cRPMI, and 3 µL samples were spotted on 7H10 agar plates supplemented with 10% v/v Middlebrook OADC. After 72 h, luminescence was measured for the second set of plates using a Synergy H4 plate reader, before recovering the cells and plating the bacteria, as done for Day 0. Agar plates were incubated for 3-4 weeks at 37°C.

### Checkerboard combination assays

The Tecan 300e Digital Dispenser was used to dispense test compounds and reference drugs in into the 96-well assay plates. *M. tuberculosis* H37Rv-LP was grown until mid-logarithmic phase (OD_590_∼0.4-0.6), inoculated to a final OD_590_ of 0.02, and incubated at 37°C for 5 days. Growth was measured by OD_590_ and data was analyzed by BLISS analysis using Combenefit software.

### Determination of intracellular ATP levels

Logarithmic phase (OD_590_∼0.4-0.6) *M. tuberculosis* was inoculated into 96-well assay plates to a final OD_590_∼0.04 and incubated at 37°C for 24 h. 50 µL BacTiter-Glo™ reagent was added to each well and the assay plate was incubated for 10 min in the dark. Luminescence was read using the Synergy H4 plate reader. Growth was determined by reading absorbance at OD_590_ after 5 days of incubation.

### Determination of oxygen consumption rate (OCR)

OCR was measured using an Agilent Seahorse XFp flux analyzer as described (18). For *M. tuberculosis*, bacterial cells were incubated in 7H9 medium supplemented with 0.01 % tyloxapol for 24 h, harvested and resuspended in unbuffered 7H9 medium without any carbon source (pH=7.35) at OD_590_∼0.4. 50 μL Agilent Cell-Tak (final concentration= 22.4 μg/mL) was added to a microtiter plate, incubated at RT for 60 min and washed with sterile water. This plate was allowed to air dry before adding 50 μL of bacterial culture. After inserting the plate in the Seahorse machine, basal OCR was measured, followed by addition of glucose. Test compounds were added and OCR was monitored for 100 min. For THP-1 cells, the cells were grown in Agilent Seahorse XF RPMI medium (pH=7.4) supplemented with 2 mM glutamine, 1 mM pyruvate and 10 mM glucose for 24h. Assay plates were coated with 50 μL Agilent Cell-Tak, 50 μL THP-1 cells were added at the density of 5×10^4^ cells/mL to each well. Basal OCR was determined, followed by addition of test compounds and monitoring changes in OCR for ∼80 min. Data was analyzed using Wave Desktop software 2.6.0.31.

### Measurement of membrane potential

Bacterial membrane potential was determined as described (19). Briefly, logarithmic phase (OD_590_∼0.4-0.6) cells were harvested and resuspended in 10 mL 7H9-Tw at OD_590_ containing 15 μM 3,3’-Diethyloxacarbocyanine iodide (DiOC2). Cells were incubated at RT for 20 min, washed, and resuspended in fresh 7H9-OADC-Tw and inoculated into 96-well assay plates to a final OD_590_ of 0.5. Assay plates were incubated at 37°C for 30 min and fluorescence measured at Ex488/Em530 (green) and Ex488/Em610 nm (red) using the Synergy H4 plate reader. The ratio of red: green fluorescence was calculated.

### Measurement of reactive oxygen species (ROS)

Intracellular ROS induction was measured as previously described (20). Briefly, logarithmic phase (OD_590_∼0.4-0.6) cells were adjusted to an OD_590_ of 1.0. 40 µM 2′,7′-dichlorofluorescin diacetate (DCFDA) was added and incubated at 37°C for 30 min. Bacteria were harvested, washed, resuspended in fresh 7H9-Tw, and inoculated into the 96-well assay plates to a final OD_590_ of 0.5. Plates were incubated at 37°C for 90 min and fluorescence read at Ex485/Em535 nm using a Synergy H4 plate reader.

### Measurement of intracellular bacterial pH

Intracellular pH was measured as previously described (21). *M. tuberculosis* expressing rGFP was grown until the logarithmic phase (OD_590_∼0.4-0.6), washed and resuspended in phosphocitrate buffer pH 4.5 (0.0896 M Na_2_HPO_4_, 0.0552 M citric acid, 0.05 % Tyloxapol). Cultures were inoculated into 96-well plates to a final OD_590_ of 0.3. Plates were incubated at 37°C for 48 h, and fluorescence read at Ex395/Em510 nm and Ex475/Em510 nm using a Synergy H4 plate reader. The ratio of Ex395/ Ex475 was calculated.

## RESULTS AND DISCUSSION

### Heterobenzamides are active against intracellular *M. tuberculosis*

We had previously identified the HBA series with promising activity against *M. tuberculosis* grown in axenic culture(9). The macrophage can serve as a niche for *M. tuberculosis* replication and forms an additional barrier for compound penetration. Therefore, we wanted to determine if our molecules could also exert activity against intracellular bacteria. We selected five representative compounds for the series (**Figure 1**): **4a** (TPN-0095204), **4m** (TPN-0099732), **4o** (TPN-0099603), **4p** (TPN-0099778) and **4zb** (TPN-0159183).

**Fig. 1.**
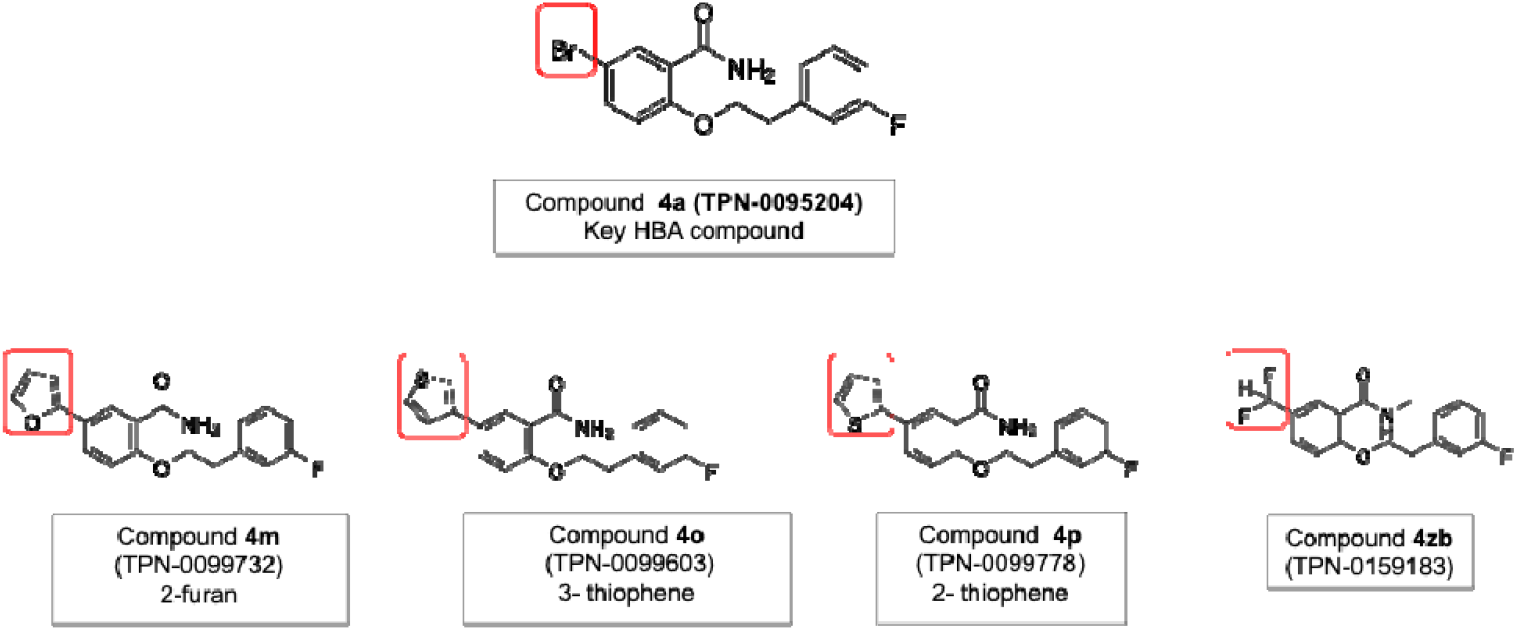
Compounds selected for this study.

We determined the IC_90_ for each compound against the wild-type strain of *M. tuberculosis* (**Table 1**). All five molecules had activity against intracellular bacteria which was comparable to the extracellular activity we had previously determined (9), confirming that molecules could enter macrophages and that the target was still vulnerable under these conditions.

**Table 1.** Activity of the compounds against extracellular and intracellular *M. tuberculosis*. ^a^IC_90_ against intracellular *M. tuberculosis*. ^b^IC_90_ against extracellular *M. tuberculosis* taken from (9). Data are the average and standard deviation of a minimum of two independent biological replicates (number of replicates in parentheses).

| Compound ID | Intracellular<sup>a</sup><br>Wild type | Extracellular<sup>b</sup> |  |  |
| --- | --- | --- | --- | --- |
|  |  | Wild type | QcrB<sub>T313I</sub> | Fold change |
| 4a | 2.6 ± 0.21 (2) | 5.5 ± 1.7 (3) | >25 (2) | >5 |
| 4m | 0.33 ± 0.18 (6) | 0.58 ± 0.34 (5) | >25 (2) | >43 |
| 4o | 0.32 ± 0.26 (5) | 0.35 ± 0.19 (3) | >25 (2) | >71 |
| 4p | 0.22 ± 0.10 (3) | 0.19 ± 0.12 (4) | >25 (2) | >132 |

### HBAs are inactive against a QcrB _**T313I**_ **mutant strain of *M. tuberculosis***

We had previously demonstrated that a QcrB_M342V_ mutant strain of M. tuberculosis demonstrated increased resistance to HBA analogs with 4-fold to >385-fold increase in IC_50_ between the two strains. We expanded our study to include the QcrB_T313I_ mutant strain, which has a mutation in the proposed binding site, and tested its sensitivity to HBA compounds (**Table 1**). We found that the mutant was highly resistant to the analogs. We also tested the activity against intracellular bacteria for the QcrB_T313I_ strain (**Figure 2**). Again, we saw a large reduction in activity against the mutant strain, where all the analogs showed greatly reduced activity against the QcrB mutant strain, although the molecules did retain some activity.

**Fig. 2.**
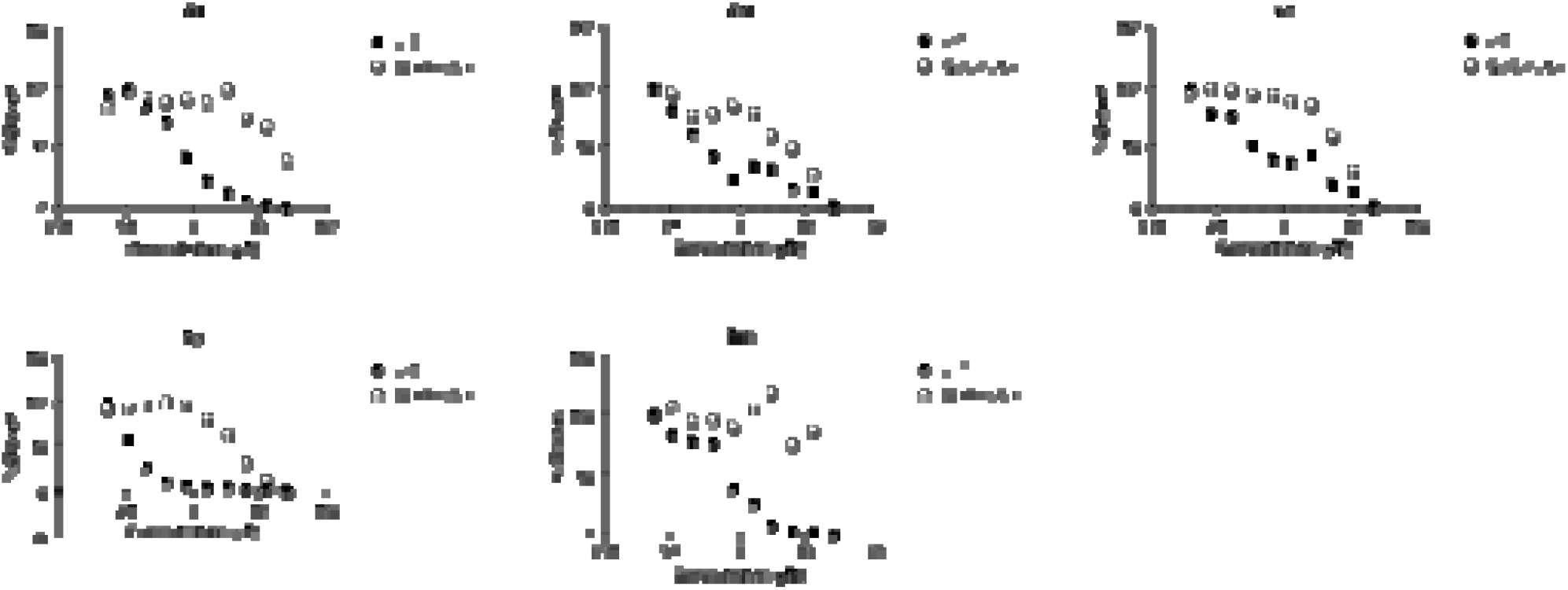
Activity against intracellular bacteria. Intracellular activity of the HBAs was determined against the wild-type and QcrB_T313I_ mutant strains. Both strains expressed luciferase constitutively, and growth was measured by luminescence after 3 days infection in THP-1 macrophage at an MOI of 1. Black circle = wild type. White circle = QcrB mutant.

### Heterobenzamides are bacteriostatic against intracellular bacteria

We and others have previously noted that QcrB inhibitors are bacteriostatic against actively replicating bacilli but are bactericidal against non-replicating bacteria (22). Since the bacteria are replicating slowly in the macrophages, we wanted to determine the kill profile for intracellular bacteria. Since our previous assay measured growth, we conducted an assay to determine if there was any kill of the original inoculum. We infected THP-1 macrophages at an MOI of 1 and incubated for 3 days. We lysed the macrophages and spotted the lysates onto agar plates to quantify viable bacteria (**Figure 3A/C**). We plated day 0 to ensure that compound carryover did not affect the recovery or growth of viable bacteria and indeed all spots grew equally. After 3 days, we saw no reduction in spot density as compared to day 3 although we able to see small differences in density when growth was inhibited. These data indicate that the molecules are inhibiting growth in the macrophage but do not kill, consistent with our previous observations that QcrB inhibitors are bacteriostatic against replicating *M. tuberculosis*. We conducted the same experiment with the QcrB_T313I_ mutant strain with similar results. No compound carryover was noted at Day 0 and no kill was obtained (**Figure 3B/D**).

**Figure 3.**
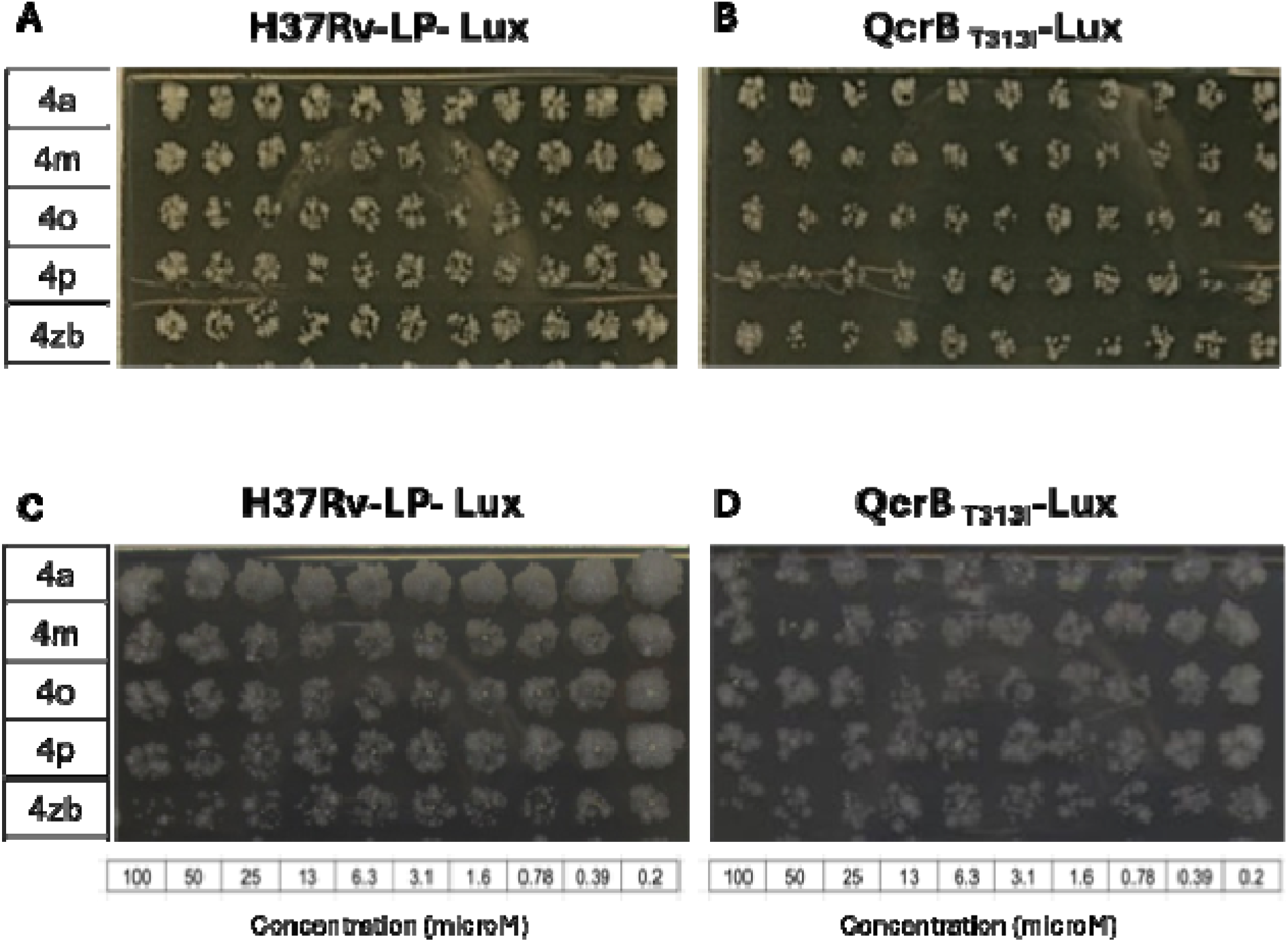
HBAs are bacteriostatic against intracellular bacteria. THP-1 cells were infected with *M. tuberculosis* strains (expressing luciferase) and incubated with two-fold serial dilutions of compounds (starting at 100 µM) for 3 days. 3 µL from each well was spotted onto agar plates. Panels A and B – Day 0. Panels C and D – Day 3. Panels A and C – wild type strain (H37Rv-LP-Lux). Panels B and D – QcrB_T313I_ mutant strain (QcrBT_313I_-Lux).

### Benzamides are active against mono-resistant strains and all *M. tuberculosis* lineages

We wanted to determine if our molecules retained activity against *M. tuberculosis* strains with resistance to the two frontline drugs, rifampicin (RIF) and isoniazid (INH), as well as determine if they had activity against different lineages of *M. tuberculosis*. We determined MICs against an INH-resistant strain (KatG_Y608*_) and a RIF-resistant strain (RpoB_S522L_). HBAs retained equivalent activity against these mono-resistant isolates (**Table 2)**. The HBAs also proved to be active against all seven geographically distinct lineages of *M. tuberculosis*. (**Table 2)**.

**Table 2.**
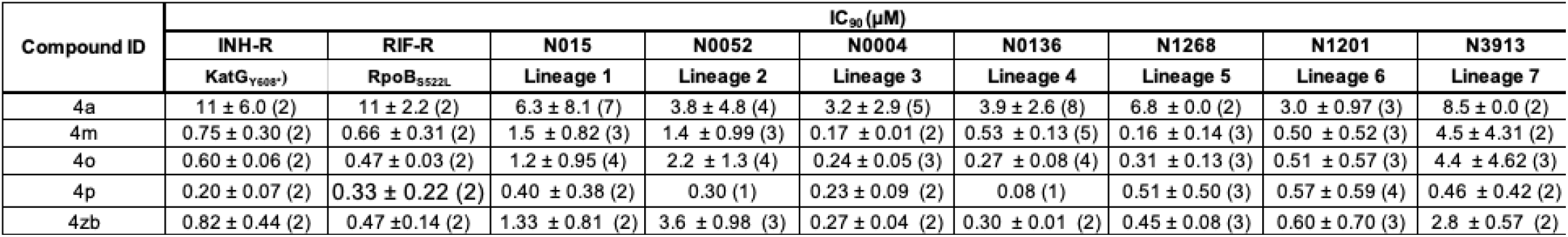
Activity of the compounds against *M. tuberculosis* strains. Data are the average and standard deviation of a minimum of two independent biological replicates (number of replicates in parentheses).

### HBAs exhibit strong synergy with bedaquiline

We investigated the interactions of the key compound **4a** with several drugs to determine if there was any synergy. We ran checkerboard assays combining **4a** with isoniazid, rifampicin, ethambutol, bedaquiline, pretomanid and linezolid (**Figure 4**). We saw strong synergy with bedaquiline and weaker synergy with isoniazid. No synergy or antagonism was seen with the other drugs.

**Figure 4.**
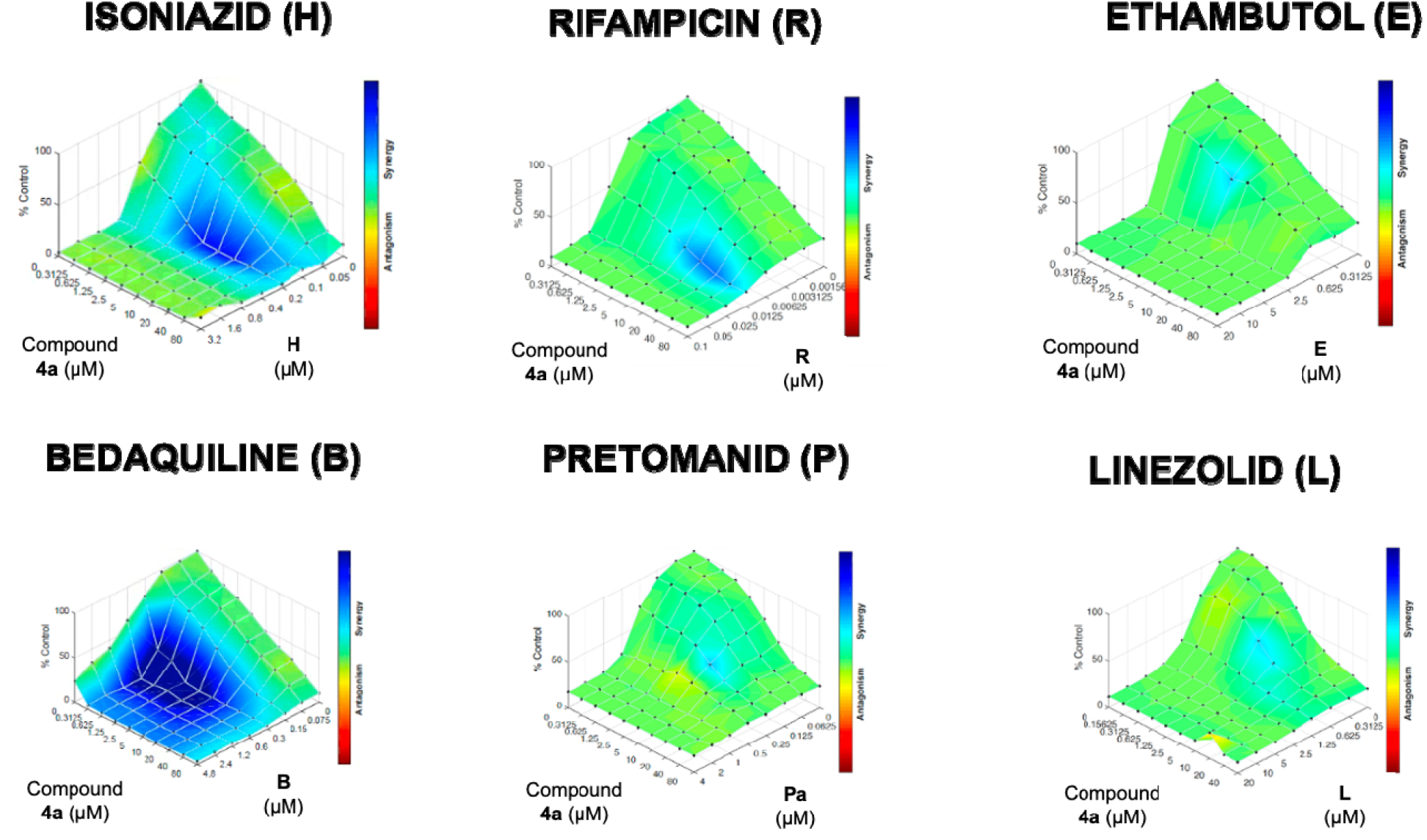
Checkerboard combination assays demonstrate synergy with bedaquiline and isoniazid. Compound **4a** was combined with each drug in a checkerboard assay. M. tuberculosis growth was measured after 5 days. Data was analyzed with Combenefit. Blue areas indicate synergy between drugs.

### HBAs causes ATP depletion and boost the oxygen consumption rate

Our data strongly suggest that HBAs target aerobic respiration by inhibiting the electron transport chain (ETC). We wanted to determine whether ATP depletion occurred, which would be expected if the ETC was perturbed. Disruption of the ETC should also lead to changes in the oxygen consumption rate (OCR) as seen with other QcrB inhibitors. We first looked at ATP levels in *M. tuberculosis* exposed to HBAs (**Figure 5**). We tested our key compound (**4a**). As expected, **4a** showed the same pattern as Q203, in that a reduction in ATP was seen and that this happened at concentrations below those which inhibited growth. Kanamycin was used as the negative control in which ATP levels correlated with growth inhibition.

**Figure 5.**
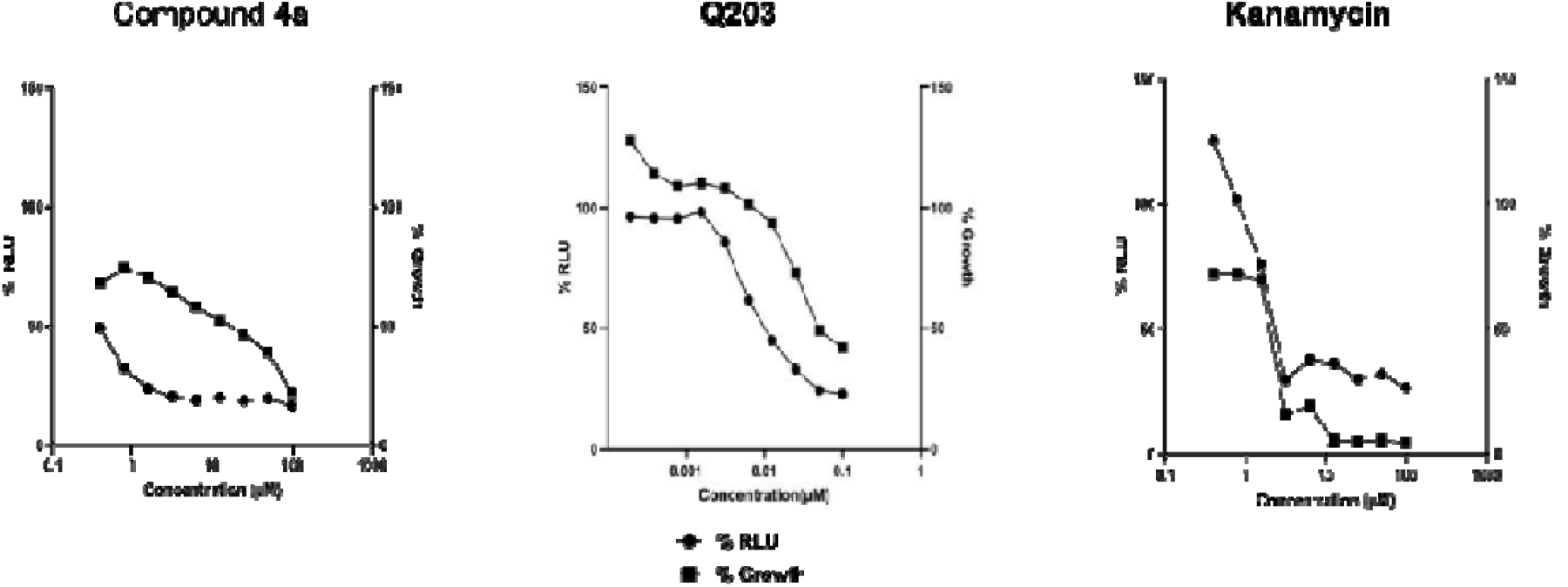
HBA 4a depletes intracellular ATP. ATP levels were measured with BacTiter-Glo after 24 h. Growth was measured by OD after 5 days. Data are normalized to the DMSO control.

ATP depletion is tightly linked with the oxygen consumption rate (OCR) thus we monitored the effect of compound **4a** on OCR in *M. tuberculosis* (**Figure 6A**). As with Q203, our compound led to a boost in OCR of similar magnitude confirming that the mode of action is via inhibition of the ETC. The potency of the HBA against the *M. tuberculosis* within THP-1 cells raised a question of the possible impact of the compound on the respiration of the THP-1 macrophages themselves. We measured the effect of compound **4a** on uninfected macrophage (**Figure 6B**). We saw no effect of our compound. Rotenone, a known inhibitor of respiration, did lead to decreased OCR in macrophages. It is interesting that QcrB inhibition leads to an increase in OCR in *M. tuberculosis* whereas inhibition of the ETC in THP-1 cells leads to a reduction. This is a well-known phenomenon in *M. tuberculosis* and has been seen with other ETC inhibitors (23).

**Figure 6.**
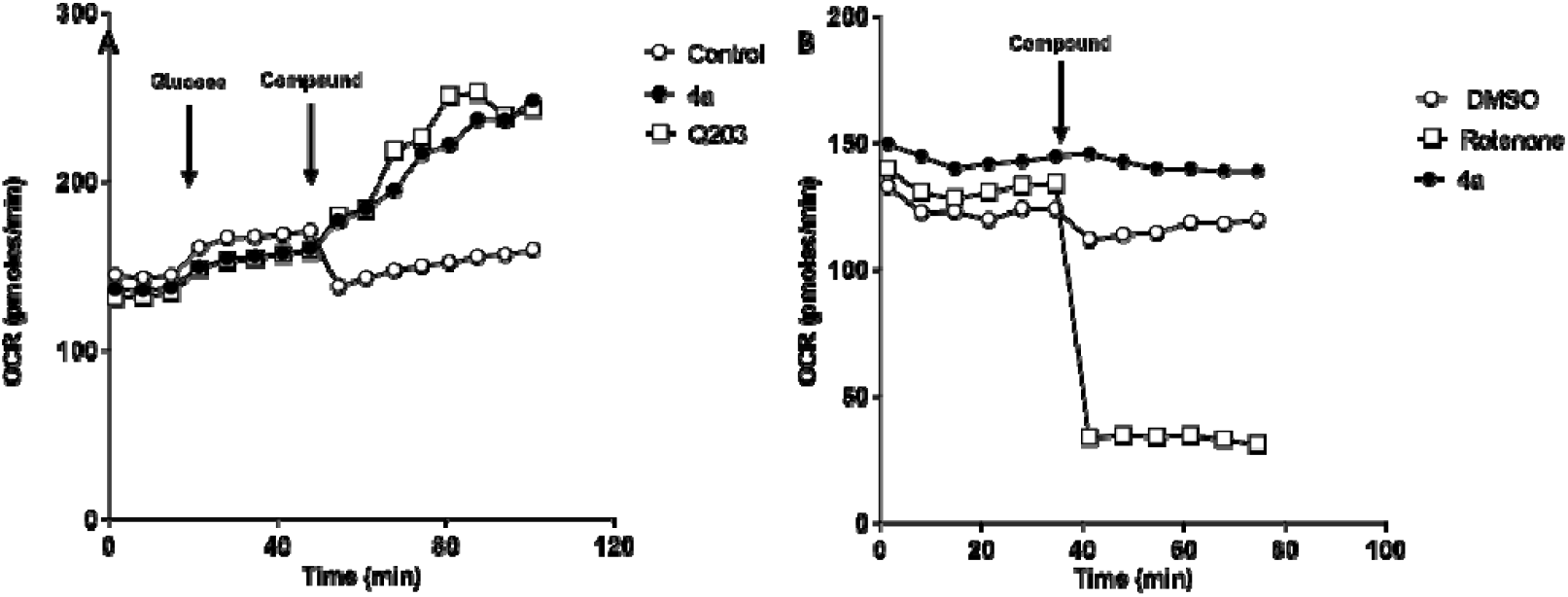
HBA 4a boosts OCR in *M. tuberculosis* but not in TP-1 cells. Oxygen consumption rate was measured using the Seahorse flux analyzer. Test compounds were added at the indicted timepoints. (A) *M. tuberculosis*. (B). THP-1 cells.

### HBA disrupted pH homeostasis, but did not induce reactive oxygen species (ROS) or alter membrane potential

We had previously noted that other inhibitors of QcrB could disrupt pH homeostasis, which is predicted since it is a proton pump and its inhibition would lead to decreased export of protons. We tested this using a strain of *M. tuberculosis* expressing a ratiometric GFP (17). We saw the same decrease in the GFP ratio for compound **4a** as for monensin, a cation ionophore (**Figure 7A**) confirming that pH homeostasis is negatively impacted. We also wanted to test whether there were other possible mechanisms of action for our series. Induction of reactive oxygen species and disruption of membrane potential are common mechanisms, so we tested for these. We saw no induction of ROS by compound **4a**, although we did see this with our positive control econazole (**Figure 7B**) (20). Similarly, we saw no perturbance of membrane potential with compound **4a**, although we were able to detect this with our positive control CCCP (**Figure 7C**). Taken together these provide further support for QcrB as the sole target of the HBA and reduce the likelihood that there are other secondary mechanisms of action.

**Figure 7.**
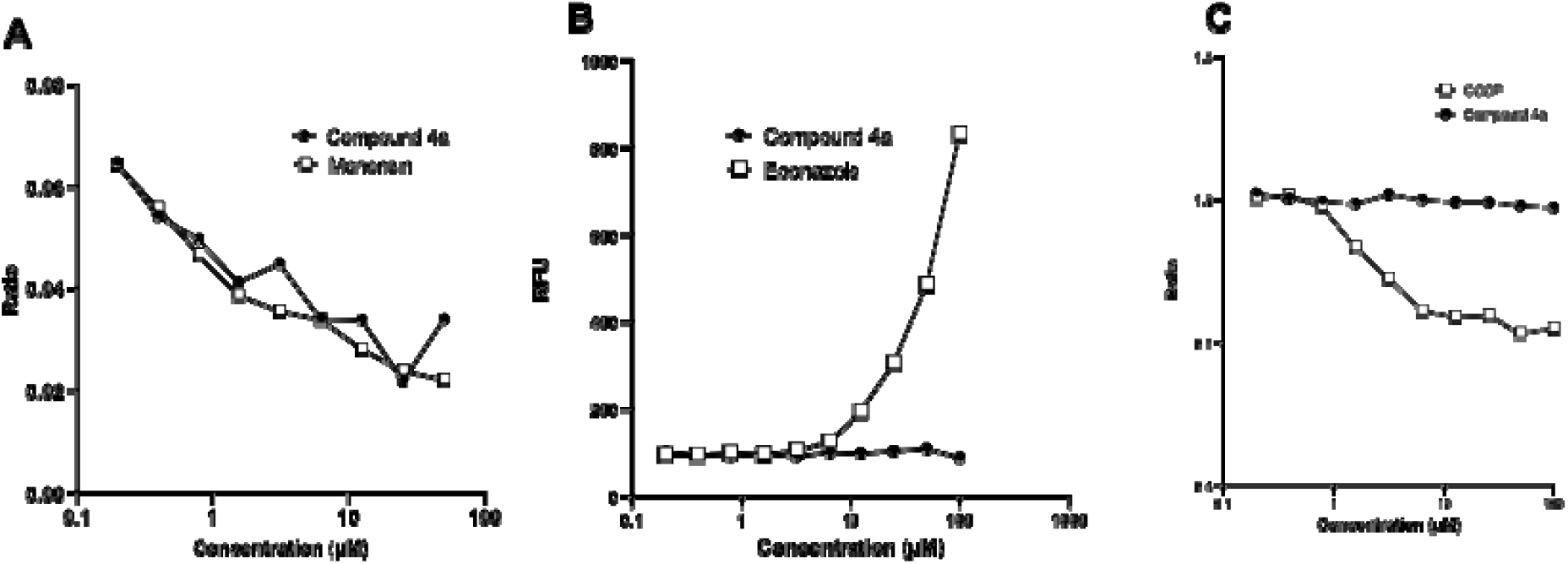
The effect of HBA compound 4a on (A) pH homeostasis (B) production of reactive oxygen species and (C) membrane potential. Control compounds were (A) monensin, (B) econazole, and (C) CCCP.

### HBAs have a narrow spectrum of activity

We wanted to determine if the HBAs were active against other bacterial species. We tested our analogs against representatives of both Gram-negative species - *Acinetobacter baumannii* ATCC 19606, *Escherichia coli* BW25113, *Enterobacter cloacae* ATCC 13047, *Klebsiella pneumoniae* ATCC 13883, *Pseudomonas aeruginosa* ATCC 10145 and Gram-positive bacteria - *Staphylococcus aureus* ATCC 12600 and *Enterococcus faecium* ATCC 19434. None of the analogs had any activity (>100 µM) (Table 3) indicating that they have a narrow spectrum of activity. The standard TB drug regimen includes multiple drugs taken over a long period of time. This runs the risk of perturbation of the healthy gut microbiota and could cause dysbiosis (24). Therefore, the narrow spectrum of the HBA species is a useful profile which would minimize the effect on the microbiome.

**Table 3.** Activity against representative Gram negative and Gram positive bacterial species. Data are from two independent biological replicates.

| Compound ID | IC<sub>90</sub> (μM) | | | | | | |
| --- | --- | --- | --- | --- | --- | --- | --- |
|  | *Acinetobacter baumannii* ATCC 19606 | *Escherichia coli* BW25113 | *Enterobacter cloacae* ATCC 13047 | *Klebsiella pneumoniae* ATCC 13883 | *Pseudomonas aeruginosa* ATCC 10145 | *Staphylococcus aureus* ATCC 12600 | *Enterococcus faecium* ATCC 19434 |
| 4a | >100 | >100 | >100 | >100 | >100 | >100 | >100 |
| 4m | >100 | >100 | >100 | >100 | >100 | >100 | >100 |
| 4o | >100 | >100 | >100 | >100 | >100 | >100 | >100 |
| 4p | >100 | >100 | >100 | >100 | >100 | >100 | >100 |
| 4zb | >100 | >100 | >100 | >100 | >100 | >100 | >100 |

### Conclusion

In this work we have further profiled the promising HBA series. The molecules are active against extracellular and intracellular bacteria with a narrow spectrum of activity. We have provided further strong evidence from multiple approaches that these molecules work by targeting QcrB, a key component of the ETC without secondary activities. In addition, and importantly for ETC inhibitors, they have no effect on macrophage respiration. Further work to develop these molecules is warranted.

## ACKNOWLEDGEMENTS

We thank Renee Allen, Lauren Ames, Arielle Butts, Sultan Chowdhury, Brock Lynde, Nikki Nguyen, Quan Pham, and Teresa Repasy for technical assistance and useful discussion.

This work was funded by the Department of Defense office of the Congressionally Directed Medical Research Programs under award number PR191269.

Seattle Children’s acknowledges that their work is conducted on the traditional, unceded land of the Coast Salish people, including the Duwamish People past and present. Seattle Children’s honors with gratitude the land itself and the Duwamish Tribe.

